# Phosphotyrosine Signal Profiling of Clinical CAR-T Reveals Tonic Signaling Associated with Therapeutic Efficacy

**DOI:** 10.1101/2024.12.22.629940

**Authors:** Bin Yao, Xueting Ye, Qian Kong, Wendong Chen, Wentao Li, Chun Feng, An He, Guoqiang Li, Lan Chen, Xiong Chen, Lina Hu, Lifen Xie, Xiao Qiu, Xi Wang, Yuhong Lin, Yixuan Cao, Jihao Zhou, Xinyou Zhang, Haopeng Wang, Ruijun Tian

## Abstract

Chimeric Antigen Receptor T (CAR-T) therapies have revolutionized the treatment of cancers such as relapsed and refractory B cell malignancies. However, the precise therapeutic mechanism of CAR-T action remain to be elucidated. In this study, we systematically analyzed CAR signaling via phosphotyrosine (pTyr) proteomics in CAR-T cells from both clinical patients and healthy donors. We found that CAR-T products from clinical patients displayed heightened tyrosine phosphorylation, particularly in the JAK-STAT, MAPK and TCR signaling cascades. We also identified the significantly regulated pTyr sites in primary CAR-T cells under tonic signaling or upon stimulation by antigen-presenting CD19-K562 cells. Although both CD28ζ and 4-1BBζ CAR-T cells exhibited comparable pTyr changes, CD28ζ CAR-T cells displayed a more pronounced activation in the TCR signaling pathway. Additionally, comparative analysis between clinical and primary CAR-T cells suggested that CAR-T products were subject to heightened tonic signaling, which may be related to therapeutic relapse. Our findings reveal the state of clinical CAR-T products and refine the CAR-T signal transduction network, providing comprehensive insights and informed guidance for CAR-T therapies.

## Introduction

CAR-T therapy involves the genetic modification of T-cells derived from a patient’s peripheral blood to incorporate a chimeric antigen receptor and then re-infusing into the patient to specifically target tumor cells^1,2^. CAR-T therapy has received much attention in recent years for its promising clinical applications in oncology^3^. Notably, it has been successfully employed into treatment for patients with relapsed and refractory B cell lymphomas, B cell acute lymphoblastic leukemia and multiple myeloma^4–6^. However, CAR-T therapies may result in serious complications such like Immune Effector Cell-Associated Neurotoxicity Syndrome (ICANS) and Cytokine Release Syndrome (CRS) and some patients still suffer from therapeutic relapse^7^.

Clinical research has extensively utilized single-cell RNA-seq transcriptomics to monitor and predict the CAR-T therapy outcomes^8–10^. Nonetheless, the correlation between RNA transcription and protein expression is typically low, highlighting the importance of multi-omics analysis in revealing signal transduction^11,12^. Additionally, several plasma or serum proteomic studies have identified potential biomarkers indicative of ICANS or CRS during CAR-T therapies^13–16^. Despite of these findings, the pattern of protein changes within clinical CAR-T cells remains poorly understood. Generally, CAR-T cells are rapidly activated upon antigen ligation, with signaling transduced through a series of protein phosphorylation, particularly tyrosine phosphorylation cascades regulated by tyrosine kinases and phosphatases. Thus, investigating post-translational modifications in CAR-T cells could provide deeper insights into the mechanisms of CAR-T therapies.

Previous studies have focused on phosphoproteomics of the different CAR-T cells using various methods^17–20^. However, several of these investigations have primarily utilized CAR-T cells derived from a immortalized cell line called Jurkat-T to compare different CAR-T signals, and pTyr profiling, particularly in the context of tonic signaling, remains insufficiently addressed. Furthermore, the number of identified pTyr sites in these studies has generally been limited and remains to be elevated. In the present study, we analyzed the pTyr proteomics of CAR-T products and peripheral blood mono-nuclear cells (PBMCs) from four clinical patients. We observed increased basal tyrosine phosphorylation in clinical CAR-T products under tonic signaling and acute pTyr changes were potentially related to therapeutic relapse. Additionally, we performed a SILAC-labeled cell-cell stimulation assay to characterize the pTyr sites participated in CAR-T signal transduction, both under tonic signaling conditions and upon activation by antigen-presenting cells. Comparison of 4-1BBζ and CD28ζ CAR-T cells revealed that both CAR-T cells exhibit similar pTyr events, with CD28ζ CAR-T cells displaying higher pTyr changes in TCR signaling pathways. Furthermore, detailed comparison between the pTyr proteomes of clinical CAR-T and primary CAR-T cells showed that clinical CAR-T cells had more extensive upregulation of pTyr levels, indicating that combination treatments might be necessary to optimize the CAR-T products and improve the effectiveness of CAR-T cell therapy.

## Results

### Identification of pTyr events in CAR-T products

To monitor the pTyr events in CAR-T products, the PBMCs and CAR-T samples were collected from four patients who were treated with second-generation anti-CD19 4-1BB-CD3z second generation CAR-T therapy cells, which have a 4-1BB-CD3z configuration. Additionally, commonly used second generation CAR-T cells, i.e., 4- 1BB-CD3z-CAR-T(4-1BBζ CAR-T) and CD28-CD3z-CAR-T(CD28ζ CAR-T), were generated using PBMCs from healthy donors^21^. To investigate changes in pTyr signaling in CAR-T cells under both tonic signaling and post-stimulation conditions, a SILAC labeled cell-cell co-culture assay was performed. pTyr peptides were enriched and analyzed via LC-MS/MS for subsequent label-free quantitative proteomic analysis (Fig. 1).

**Fig. 1.**
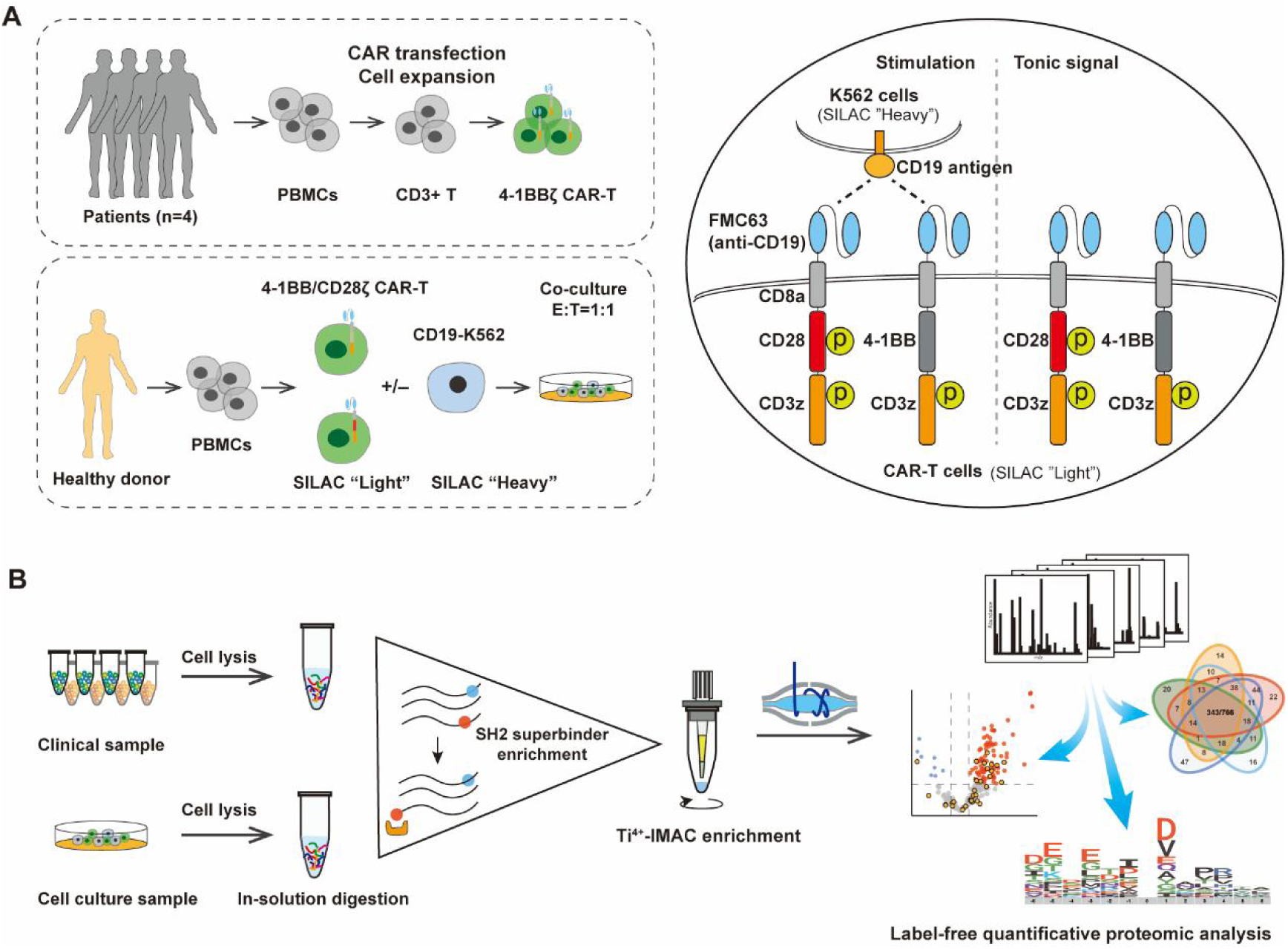
MS-based phosphoproteomic analysis of PBMCs and CAR-T cells. A. Schematic of work flow of tyrosine phosphoproteomic profiling including CAR-T cells and PBMCs from clinical patients and healthy donors. CD28ζ and 4-1BBζ CAR-T cells were stimulated with/without CD19-K562 cells. B. Cells were lysed and digested, and the tyrosine-phosphorylated peptides were enriched by the SH2 superbinder protein and Ti4^+^-IMAC beads and then analyzed by LC-MS/MS.

Given the inherent challenges in identifying pTyr sites, we initially conducted an analysis of protein levels and phosphorylation sites in PBMC samples. This analysis identified over 5000 protein groups in both CD3+ and CD3− populations isolated from PBMCs of clinical patients, as well as in CAR-T cells derived from healthy donors (Fig. 2A, B). We also identified more than 6000 phosphorylation sites across both CD3+ and CD3- populations isolated from three clinical PBMC samples. Among these phosphorylation sites, 88.97% (7879) were phosphoserine (pSer), 9.27% (821) were phosphothreonine (pThr), and 1.76% (156) were pTyr (Fig. 2C). Following pTyr-specific enrichment, a total of 363 pTyr sites were identified in CAR-T samples from patients, and 766 pTyr sites were identified in CAR-T cells derived from healthy donors (Fig. 2D). In summary, the identification of these pTyr sites (a total of 1028 pTyr sites) can offer valuable insights into the phosphorylation events and signaling mechanisms that regulate CAR-T cell function.

**Fig. 2.**
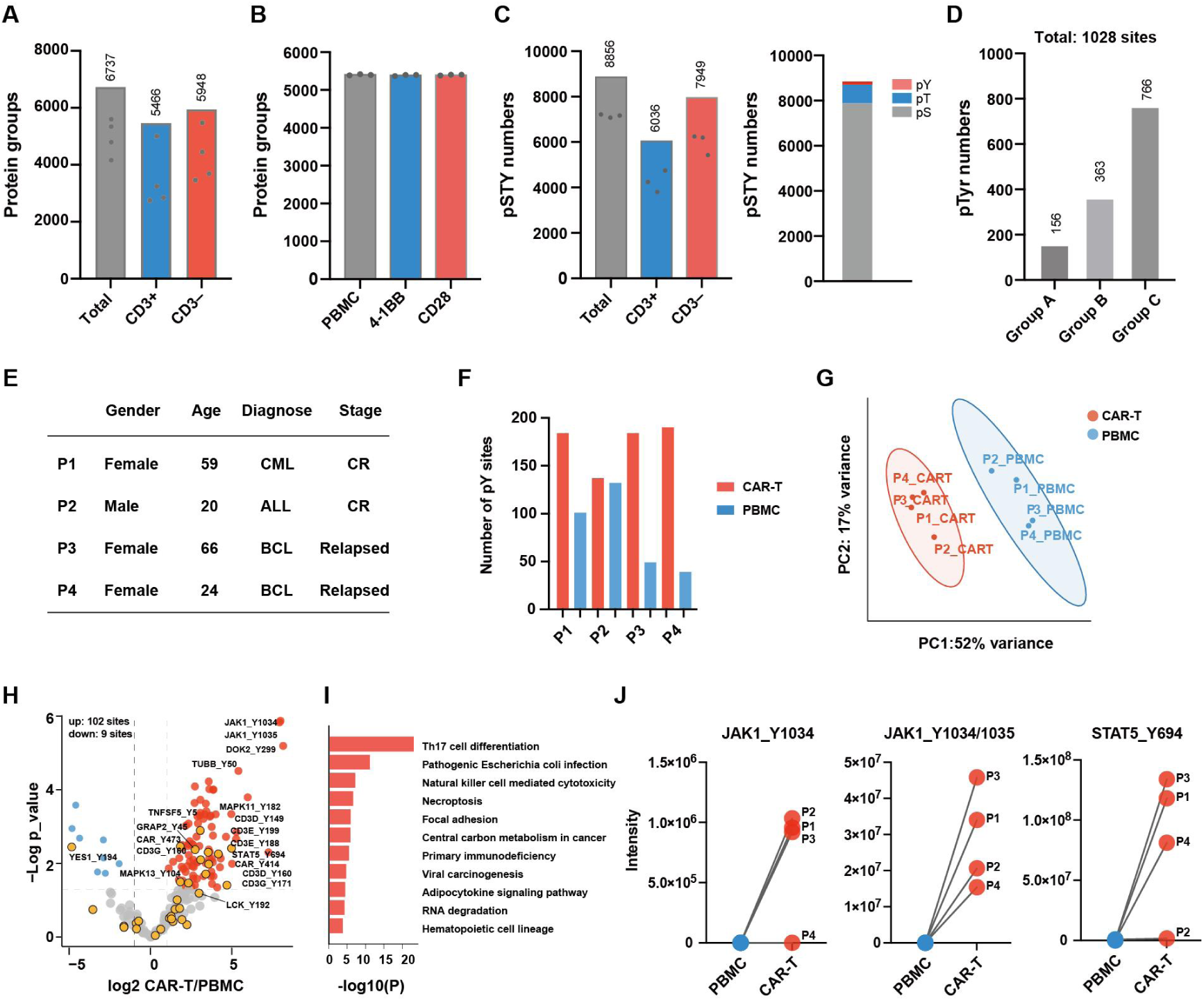
pTyr profiling of clinical CAR-T cells. A. Number of proteins identified in PBMCs from clinical patients (n = 4), with the total number of proteins shown above. B. Number of proteins identified in primary CAR-T cells derived from healthy donors. C. Number of phosphorylated sites identified in PBMCs from patients (n = 3), with the total number of pSer, pThr, and pTyr sites shown in the right panel. D. Total number of pTyr sites identified across different experimental groups. (group A, pTyr sites identified in PBMCs from clinical patients, as shown in panel C; group B, pTyr sites identified in clinical CAR-T cell samples; group C, pTyr sites identified in primary CAR-T cells from healthy donors. In groups B and C, pTyr peptides were enriched using SH2 superbinder protein.) E. Clinical information of the four patients undergoing CAR-T therapy. CML: chronic myelogenous leukemia; ALL: Acute lymphoblastic leukemia; BCL: B cell lymphoma; CR: complete response. F. Number of identified pTyr sites in PBMCs and CAR-T cell products. G. Principle component analysis of pTyr profiles in CAR-T and PBMC cells. H. Volcano plot comparing clinical CAR-T cells to PBMCs. Significantly up-regulated and down-regulated pTyr sites are labeled in red and blue respectively. Sites in the T cell receptor signaling pathway are highlighted in yellow dots. Significance threshold: S0=1, FDR<0.05. I. KEGG pathway enrichment analysis of significantly up-regulated pTyr sites from panel H. J. Intensity plots of selected up-regulated pTyr sites in the JAK-STAT signaling pathway from panel H.

### Clinical CAR-T products exhibit high tonic signaling

We first analyzed the pTyr data from clinical samples and identified between 40 and 170 pTyr sites across different samples (Fig. 2E and F). Principal component analysis (PCA) revealed distinct patterns of pTyr site distribution between PBMCs and CAR-T cells, indicating significant changes in the pTyr proteome after CAR transduction (Fig. 2G). Additionally, the volcano plot comparing CAR-T cells to PBMCs showed more up-regulated pTyr sites compared to down-regulated ones. Notably, several pTyr sites related to the T cell receptor (TCR) signaling pathway were up-regulated in CAR-T cells (Fig. 2H). KEGG pathway enrichment analysis showed significant enrichment in Th17 cell differentiation and Natural killer (NK) cell-mediated cytotoxicity pathways (Fig. 2I). Furthermore, phosphorylation at JAK _Y1034/Y1035 and STAT5_Y694, both of which play critical roles in cell proliferation and differentiation, was markedly elevated, suggesting that clinical CAR-T products may undergo self-activation^19,22–24^ (Fig. 2J).

To have an overview of pTyr changes, pTyr sites were clustered in a heatmap (Fig. S1A). Interestingly, in the first cluster, pTyr sites, including those at CD247 (CD3z in CAR) in patients P3 and P4, were significantly up-regulated in CAR-T cells compared to PBMCs. Although CAR_Y414 and CAR_Y473 were significantly up-regulated in volcano plot, their intensity changes varied (Fig. S1B). The higher phosphorylation of tyrosine in CAR in P3 and P4 suggests increased activation of these CAR-T products under tonic signaling. Furthermore, the global phosphorylated tyrosine changes were more pronounced in P3 and P4, despite similar median of the log2 fold changes across all four patients. Moreover, higher median pTyr in TCR signaling pathway in P3 and P4 indicates that a potential link between the hyper-activation of CAR-T products and the efficiency of CAR-T therapy (Fig. S1C).

To further evaluate this conclusion, another CAR-T product was collected from a patient with acute lymphoblastic leukemia (ALL). pTyr profiling showed a great number of phosphorylated tyrosine sites in CAR-T sample compared to PBMCs, with a significant up-regulation in tyrosine phosphorylation (Fig. S1D, E). Notably, the pTyr sites in the TCR signaling pathway were up-regulated, particularly at CAR proteins, indicating activation of this CAR-T product (Fig. S1F). Furthermore, comparison of the fold change in pTyr sites between PBMCs and CAR-T cells revealed significantly higher levels in samples from relapsed patients. This suggests that alterations in tyrosine phosphorylation could serve as potential biomarkers for predicting therapeutic outcomes in clinical CAR-T therapy (Fig. S1G).

### Profiling of pTyr proteome of primary CAR-T cells

To better understand the signal transduction mechanism in CAR-T cells, we constructed two widely used 2^nd^ generations CARs types, i.e., 4-1BBζ and CD28ζ CARs. The differences between 4-1BB and CD28 co-stimulatory domain were depicted, with tyrosine sites highlighted in red^25,26^ (Fig. 3A). As indicated earlier, to investigate the CAR-T signal transduction in the presence or absence of antigen-presenting cancer cells, we employed SILAC method to label CAR-T cells and CD19-K562 cancer cells prior to co-culture assay. Following a 5-minute stimulation, we enriched the pTyr peptides and performed LC-MS/MS analysis (Fig. 3B). A total of 766 pTyr sites were identified, with approximately 300-400 sites identified in each replicate (Fig. 3C and S1D). The label-free quantification (LFQ) intensities of identified tyrosine phosphorylated peptides showed high correlations (Fig. 3D). Venn diagram showed that 343 of a total of 766 pTyr sites overlapped among five groups (Fig. 3E). Moreover, PCA analysis indicated the difference in the pTyr profiles among the experimental groups, suggesting that CAR-T cells exhibit distinct signaling characteristics upon activation (Fig. 3F).

**Fig. 3.**
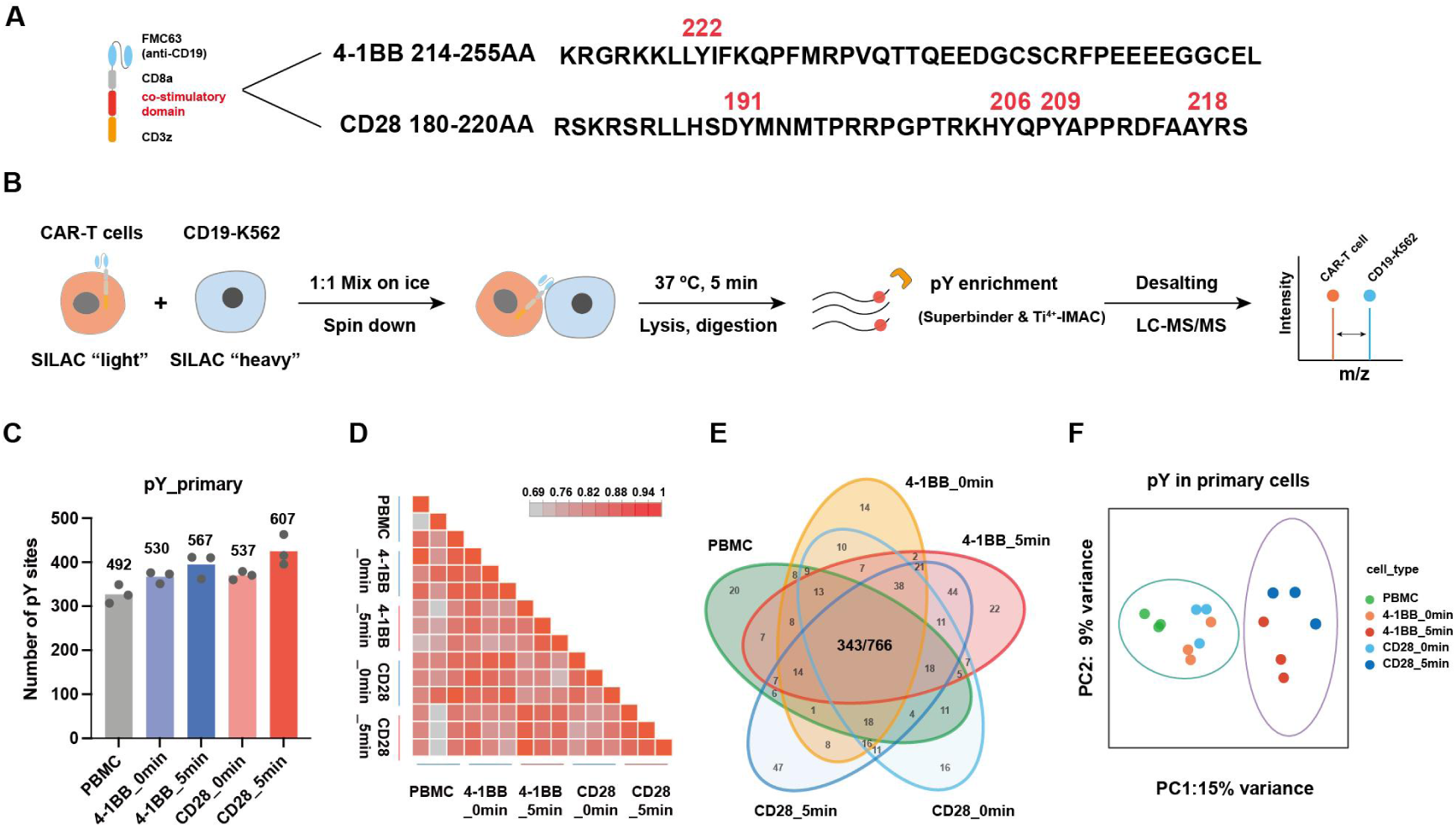
pTyr profiling of CAR-T upon stimulation by antigen-presenting cells. A. Schematic representation of CAR construction. The amino acids of 4-1BB or CD28 co-stimulatory domains are shown with tyrosine highlighted in red. B. Experimental design for comparing different CAR-T signaling. C. Number of identified pTyr sites across different experiments, with the total number of identified pTyr sites in each group labeled above. D. Correlation plot of the pTyr intensities from different experiments. E. Venn diagram showing the overlap of pTyr sites among different groups. F. Principle component analysis of pTyr profiles across different groups.

### 4-1BBζ and CD28ζ CAR-T cells show similarities under tonic signaling

To investigate the pTyr events under tonic signaling, we first compared the significantly changed pTyr sites in unstimulated 4-1BBζ or CD28ζ CAR-T cells. Volcano plots showed 17 significantly up-regulated and 4 down-regulated pTyr sites in 4-1BBζ CAR-T cells, and 20 and 11 pTyr sites showed significantly up-or down-regulated in CD28ζ CAR-T cells, respectively (Fig. 4A). Several pTyr sites exhibited similar changes, including HSP90AA1_Y492, LCP_Y39 and MEMO_Y210 (Fig. 4B). The heat shock protein HSP90AA1 is a chaperone that plays important roles in cell proliferation^27^. MEMO has been implicated in facilitating cell migration and polarity^28,29^, while LCP2, also known as SLP-76, is an adaptor protein in TCR signaling pathway that mediates signaling following T cell activation^30,31^. Phosphorylation at these sites may impact the functions of these proteins although this has not been previously reported. In addition, up-regulated pTyr sites in NEDD9 proteins, (Y92 and Y166 in CD28ζ CAR-T and Y317 in 4-1BBζ CAR-T) have been shown to be phosphorylated by the ABL kinase family, which were essential for T-cell migration^32^. In summary, these findings suggests that CAR-T cells are self-activated with considerably enhanced cell mobility.

**Fig. 4.**
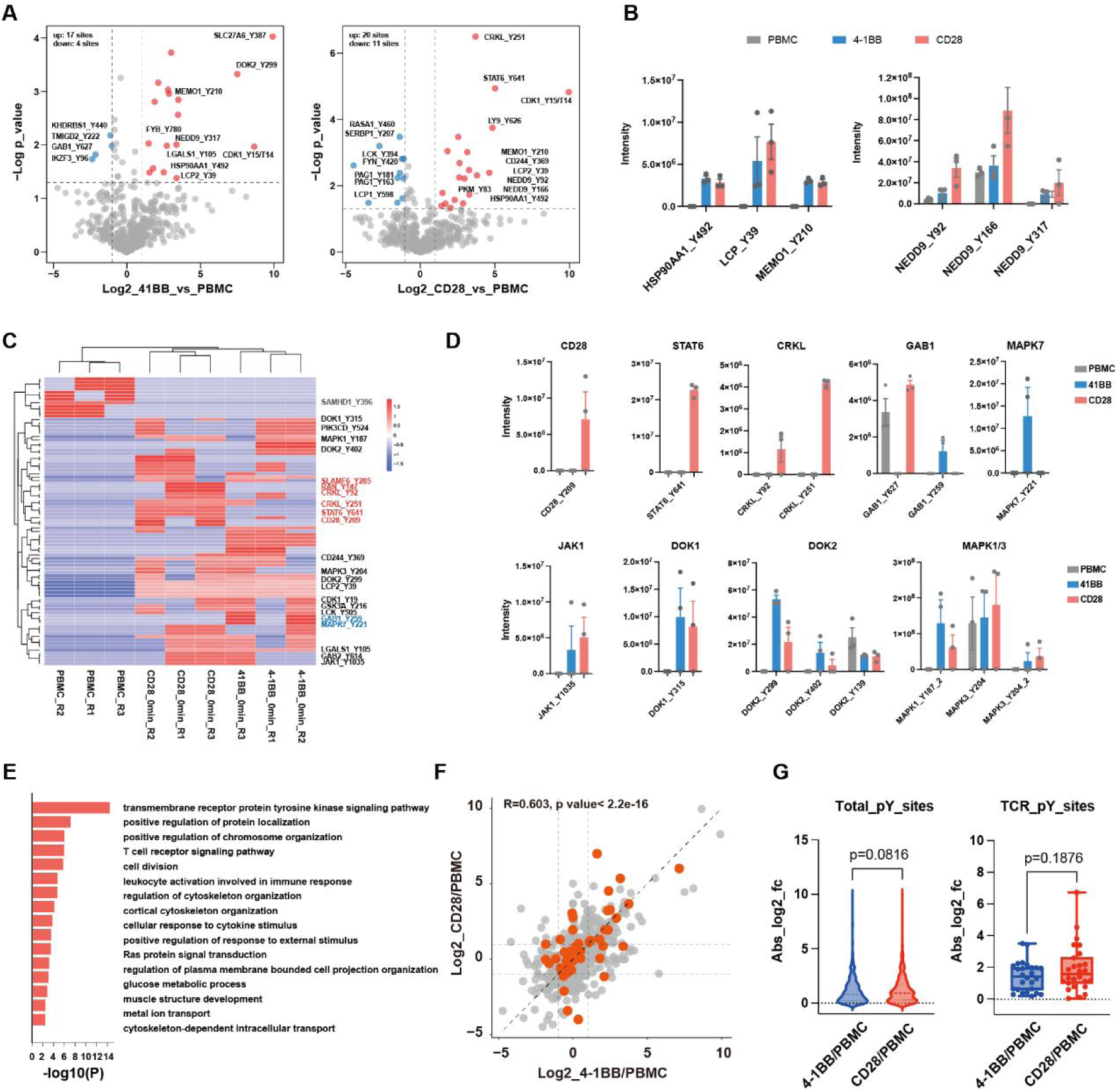
4-1BBζ and CD28ζ CAR-T exhibit similar tonic signaling. A. Volcano plots showing the significantly regulated pTyr sites in unstimulated 4-1BBζ or CD28ζ CAR-T cells versus PBMCs. Significantly up-and down-regulated pTyr sites are shown in red and blue, respectively. (cutoff: Log2 fold change > 1 or Log2 fold change ˂-1 and-log 10 p_value>1.3). B. Intensities of selected significantly altered pTyr sites from panel A. C. Heatmap highlighting unique pTyr sites in PBMCs or CAR-T cells. 4-1BBζ or CD28ζ CAR-T unique pTyr sites are marked in blue and red, respectively. D. Intensities of selected significantly altered pTyr sites from panel C. E. KEGG pathway enrichment analysis of shared unique pTyr sites identified from panel C. F. Comparative analysis of the log2 in 4-1BBζ CAR-T/PBMCs and log2 CD28ζ CAR-T/PBMCs. Red dots represent pTyr sites within the TCR signaling pathway. G. Absolute log2 fold change of CAR-T cells/PBMCs for total pTyr sites (left panel) and the pTyr sites identified within the TCR signaling pathway (right panel).

We then identified pTyr sites unique to 4-1BBζ or/and CD28ζ CAR-T cells. Consequently, the filtered pTyr sites showed 12 unique pTyr sites in the PBMCs group, and 11 and 10 unique sites in unstimulated 4-1BBζ or CD28ζ CAR-T cells, respectively. Specifically, CD28_Y209, STAT6_Y641, CRKL_Y92 and CRKL_Y251 were identified in CD28ζ CAR-T cells under tonic signaling. The phosphorylation of MAPK7_Y221 (ERK5) was up-regulated in 4-1BBζ CAR-T cells, while GAB1_Y259 and GAB1_Y627 showed differential phosphorylation between these two CAR-T cells. Additionally, 52 pTyr sites, e.g., JAK1_Y1035, DOK1_Y315, DOK2_Y299 and Y402, MAPK1_Y187 (ERK2) and MAPK3_Y204 (ERK1), show higher phosphorylation in both unstimulated 4-1BBζ and CD28ζ CAR-T cells, indicating shared phosphorylation events in these two CAR-T cells (Fig. 4C, D). In particular, phosphorylation at JAK1_Y1035 and DOK2_Y299 was also up-regulated in clinical CAR-T cells, confirming that these phosphorylation changes occur under tonic signaling.

We next analyzed the shared pTyr sites in 4-1BBζ and CD28ζ CAR-T cells with Gene Ontology Biological Process (GOBP) enrichment. As a consequence, transmembrane receptor protein tyrosine kinase signaling pathway and T cell receptor signaling pathway were highlighted as top enriched pathways, suggesting the activation of CAR-T cells (Fig. 4E). The Log2 fold changes of identified pTyr sites in 4-1BBζ and CD28ζ CAR-T cells under tonic signaling were similar, although that tyrosine in TCR pathway proteins exhibiting slightly higher phosphorylation in CD28ζ CAR-T cells (Fig. 4F, G). Previously studies have shown that the anti-CD19 4-1BBζ and CD28ζ CAR-T cells have similar global transcriptional profiles under tonic signaling^33^, supporting our findings that these two CAR-T cell types are highly similar in this context.

### CD28**ζ** CAR-T cells exhibit enhanced activation at TCR signaling pathway

To interrogate the pTyr proteome changes after CAR-T stimulation, we compared the pTyr levels between stimulated and unstimulated 4-1BBζ or CD28ζ CAR-T cells. Significant regulated pTyr sites were observed in both stimulated CAR-T cells, with 47 and 54 sites up-regulated and 17 and 11 down-regulated in 4-1BBζ and CD28ζ CAR-T cells, respectively (Fig. 5A). Specifically, certain tyrosine sites, such as Y292, Y492 and Y493 on ZAP70, Y579 and Y580 on PTK2B, Y1253 on PLCG1, Y187 on MAPK1, and Y204 on MAPK3, as well as several tyrosine sites on p38 MAPKs, were previously known to be phosphorylated following T cell activation^34–39^. These results indicate that the signal transduction in both 4-1BBζ and CD28ζ CAR-T cells is successfully activated after co-culture with antigen-presenting CD19-K562 cells at 5 minutes. KEGG pathway enrichment analysis revealed that both CAR-T cell types activated similar pathways upon stimulation. However, CD28ζ CAR-T cells exhibited significantly greater enrichment in TCR signaling and NK cell-mediated cytotoxicity pathways, which is consistent with the enhanced anti-tumor effects and NK-like cytotoxic activity observed in CD28ζ CAR-T cells^21^ (Fig. 5B, Fig. S2A).

**Fig. 5.**
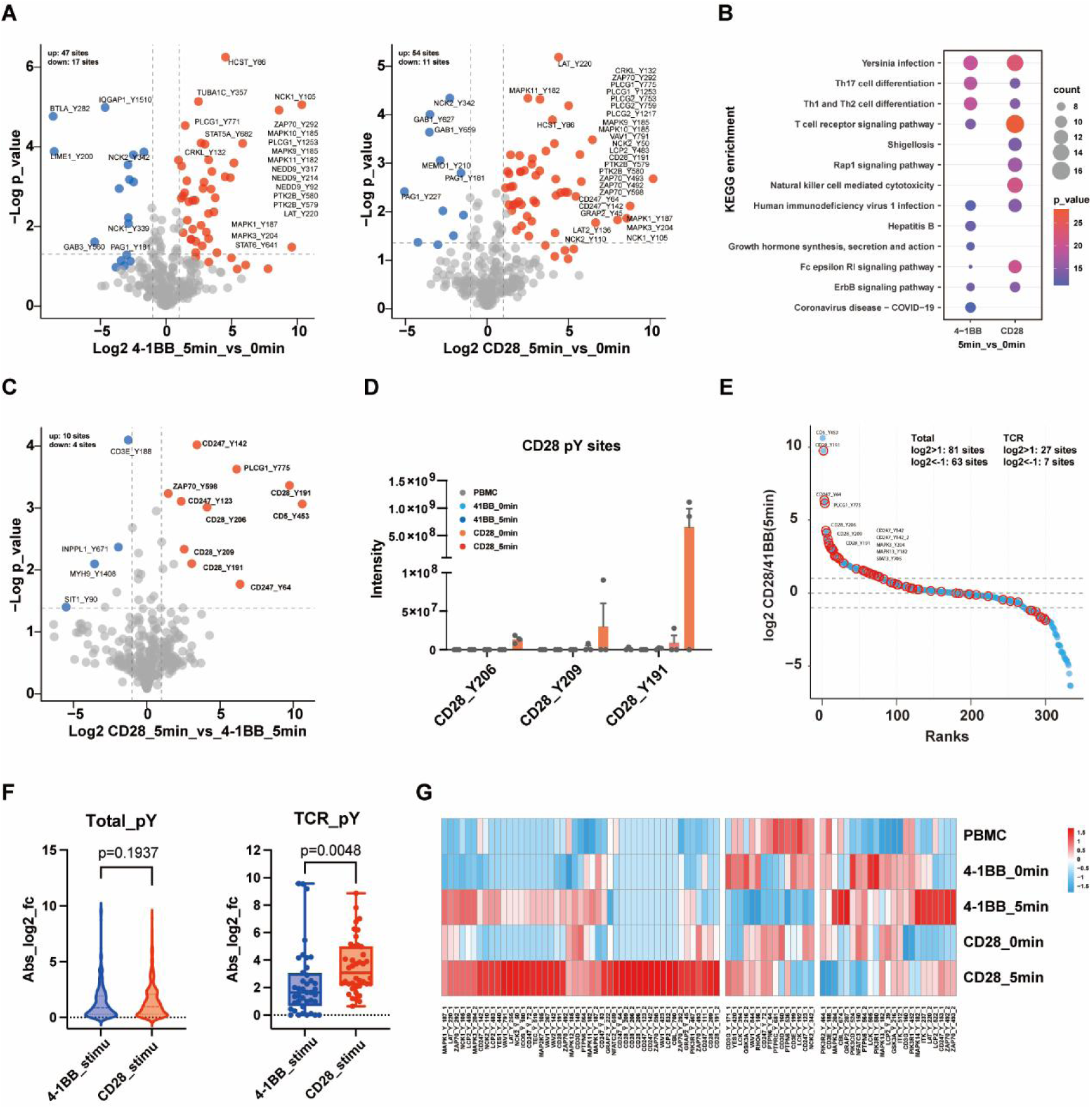
CD28ζ CAR-T cells show stronger TCR signal upon stimulation compared to 4-1BB. A. Volcano plot illustrating the differential pTyr changes in 4-1BBζ or CD28ζ CAR-T post-stimulation by CD19-K562 antigen-presenting cells. Significantly up-and down-regulated pTyr sites are shown in red and blue, respectively. Significance threshold: S0=1, FDR<0.05. B. KEGG pathway enrichment analysis of significantly up-regulated pTyr sites in stimulated CAR-T cells, presenting with the top10 enriched pathways in each dataset. C. Volcano plot showing differential pTyr changes in stimulated CD28ζ versus 4-1BBζ CAR-T cells. Significantly up-and down-regulated pTyr sites are shown in red and blue, respectively. Significance threshold: S0=1, FDR<0.05. D. Intensities of pTyr sites within CD28ζ co-stimulatory domain. E. pTyr sites ranked by the log2 fold change in stimulated CD28ζ versus 4-1BBζ CAR-T cells. pTyr sites in the TCR signaling pathway are highlighted in red circle. F. Absolute fold change of identified pTyr sites in CAR-T cells under tonic signaling or post-stimulation by CD19-K562 cells. G. Heatmap displaying the z-score of intensities of the pTyr sites across different groups.

We also compared the frequency of amino acids types in the identified up-regulated peptide sequences. The phosphorylated sequence in both 4-1BBζ and CD28ζ CAR-T cells showed similarity, with D or E from-3 to-1 position, D/V/E/Q at position +1, and hydrophobic amino acids at position +3, suggesting motifs related to kinase/phosphatase substrate and SH2 domain binding motifs^40–43^ (Fig. S2B). Moreover, protein domain enrichment analysis also revealed significant enrichment of the SH2, SH3, and kinase-related domains in both 4-1BBζ and CD28ζ CAR-T cells after stimulation (Fig. S2C). Kinase-substrate analysis (KSEA) indicated significant activation of JAK2, JAK3, MAP2K2, MAP2K1, MAPK3, and ZAP70 kinase in stimulated CAR-T cells, highlighting the activation of the JAK-STAT, MAPK and TCR signaling pathways, corroborated by KEGG pathway enrichment analysis (Fig. S2D).

Previous reports have shown that CD28ζ CAR-T cells display stronger T cell activation signals than 4-1BBζ CAR-T cells^18^. To this end, we compared the pTyr proteome in stimulated CD28ζ and 4-1BBζ CAR-T cells. It is well-known that the difference between CD28ζ and 4-1BBζ CAR lies in the co-stimulatory domain, with several tyrosine sites in CD28ζ can be phosphorylated post-stimulation. As expected, pTyr sites in CD28ζ co-stimulatory domain were remarkably up-regulated in stimulated CD28ζ CAR-T cells. Meanwhile, pTyr sites in CAR, PLCG1, and ZAP70 were also up-regulated in CD28ζ CAR-T cells, indicating a higher level of downstream signal of tyrosine phosphorylation (Fig. 5C, D). Furthermore, by mapping pTyr sites in TCR signaling pathway proteins onto a scatter plot, we observed that the majority of these sites were up-regulated in stimulated CD28ζ CAR-T cells compared to 4-1BBζ CAR-T cells, while the global pTyr sites showed nearly identical distributions between up-and down-regulated sites (Fig. 5E). Consistently, the log2 fold changes of pTyr sites in TCR signaling pathway proteins were significantly higher in CD28ζ CAR-T cells upon stimulation (Fig. 5F). As detailed in heatmap, the tyrosine sites which showed higher phosphorylation were in CD28, CD247(CD3z in CAR), ZAP70, VAV1, PIK3R1, MAPK1, and MAPK3 proteins, indicating stronger activation in TCR signaling pathway for CD28ζ CAR-T cells (Fig. 5G).

To comprehensively analyze the pTyr changes, we performed a time-course clustering analysis. As a consequence, from a total of 12 clusters, the top 4 clusters of particular interest were those that represented pTyr sites up-regulated in all CAR-T cells (cluster 3), sites up-regulated in stimulated CAR-T cells (cluster 7), sites down-regulated in stimulated CAR-T cells (cluster 5) and sites more abundant in stimulated CD28ζ CAR-T cells (cluster 12), respectively. KEGG pathway enrichment analysis showed that the most significant pathway enriched in these clusters was related to T cell activation. Specifically, cluster 7 included sustained up-regulation of JAK1_Y1034 and STAT5A_Y90, indicating the proliferation of CAR-T cells (Fig. S3A). Cluster 3 showed up-regulation of PTK2B_Y579/Y580 and MAPK1_Y187 in stimulated CAR-T cells, while cluster 5 showed down-regulation of CD3E_Y199 and PTPN11_580, illustrating activation of CAR-T cells by antigen-presenting cells, aligning with previous studies^36,37,44–46^ (Fig. S3B, C). Additionally, phosphorylation on ZAP70_Y493 and NFATC2_Y659 were more up-regulated in stimulated CD28ζ CAR-T cells compared to 4-1BBζ CAR-T cells, which may also explain the stronger tumor-killing abilities of CD28ζ CAR-T cells (Fig. S3D).

### Clinical CAR-T cells show overall up-regulated tyrosine phosphorylation

As mentioned above, the clinical CAR-T cells displayed an active phenotype, with the majority of pTyr sites being up-regulated. Notably, the volcano plots of tonic and stimulation signals of CAR-T cells from healthy donor showed more symmetrical distributions compared to those from clinical CAR-T cells (Fig. 2H, 4A and 5A). We then compared the up-regulated pTyr sites in clinical samples with those up-or down-regulated pTyr sites in 4-1BBζ or CD28ζ CAR-T cells from healthy donors. The number of overlapped sites between clinical and tonic/stimulation conditions were nearly identical. For instance, pTyr sites such as PLCG1_Y783, DOK2_Y299, SLAMF1_Y281, HCST_Y86 (PIK3AP_Y86), and PRKCD_Y313, among others, showed up-regulation upon stimulation, whereas these pTyr sites were already up-regulated in unstimulated clinical CAR-T products^47–50^ (Fig. 6A and S4). Furthermore, some pTyr sites exhibited distinct regulatory patterns, which can be identified by overlapping the up-regulated sites in clinical CAR-T products with down-regulated sites in primary CAR-T cells upon stimulation. For example, phosphorylation at Y199 in CD3E, Y542 and Y580 in PTPN11, and Y227 and Y359 in PAG1 were down-regulated after CAR-T stimulation, while these sites were regulated conversely in clinical CAR-T cells (Fig. 6B). In principle, these sites or proteins have been shown to negatively regulate T cell signaling following activation^51–53^. This suggested that these clinical CAR-T cells may be in a hyper-activated state and may require modulations. In summary, the results demonstrated that the clinical CAR-T products exhibit unique features compared to primary CAR-T cell lines, underscoring the needs for monitoring the activation state of clinical CAR-T products and pharmacological interventions.

**Fig. 6.**
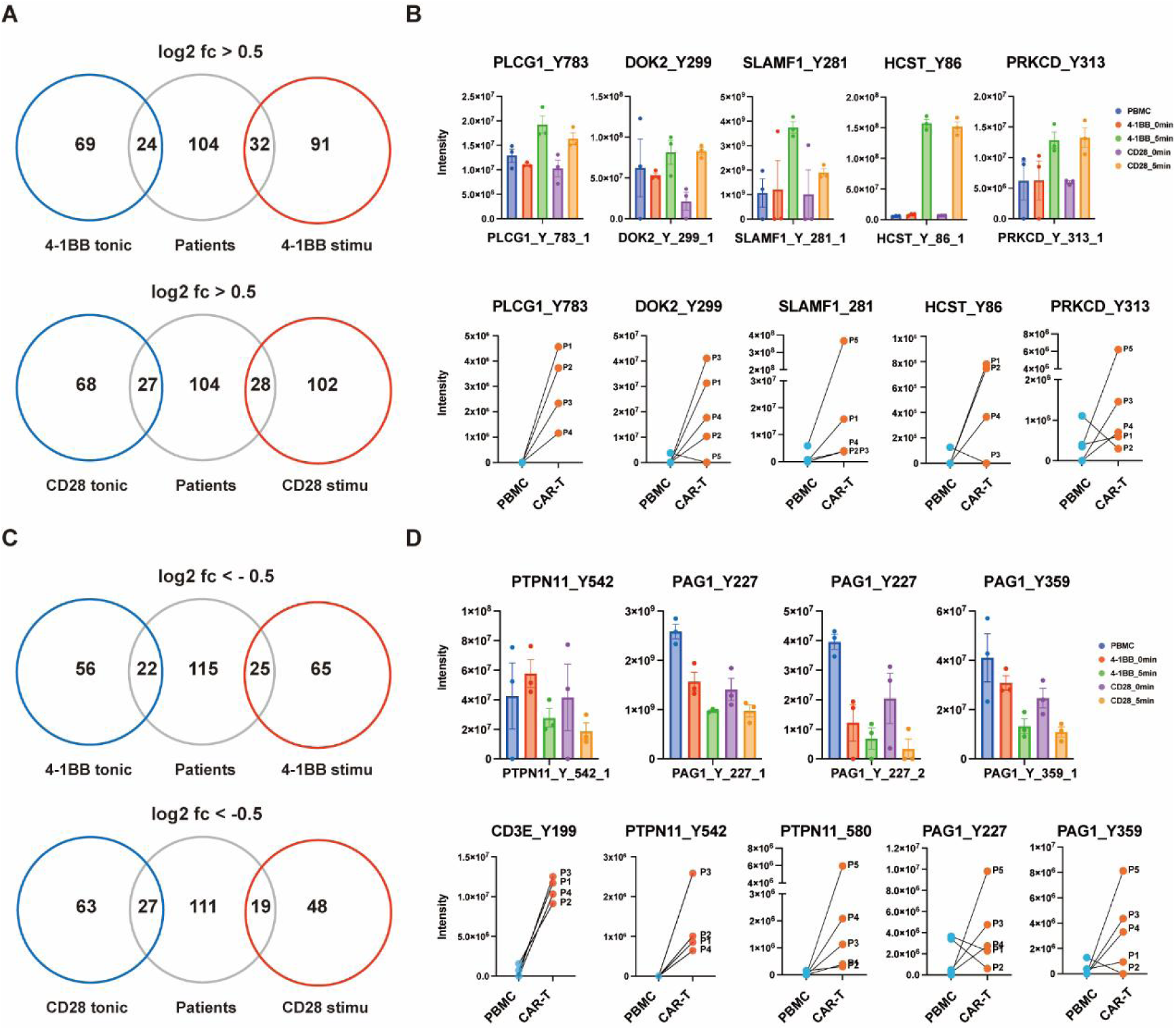
Clinical CAR-T products show different feature in tonic signaling. A. Venn diagram showing the overlap of up-regulated pTyr sites across different datasets. B. Intensity plots of selected pTyr sites in panel A. C. Venn diagram showing the overlap between up-regulated pTyr sites in clinical CAR-T and down-regulated pTyr sites in primary CAR-T cells. D. Intensity plots of selected pTyr sites identified in panel C.

## Discussion

CAR-T therapy has received remarkable success in treating cancers, particularly in patients with relapsed or refractory B cell malignancies. Despite these advances, challenges such as CRS and ICANS persist, and some patients experience relapse during therapies. Moreover, standardized manufacturing quality management for clinical CAR-T products remains inadequate. Previous studies have primarily focused on the transcriptomics or serum proteomics to identify differential T cell subsets and chemokine production of CAR-T cells, as well as to predict the therapeutic outcomes. However, post-translational modifications, especially the pTyr in CAR-T products and during the CAR-T proliferation post-infusion, have received insufficient attention. Furthermore, existing studies on pTyr proteomics of tonic and activated CAR-T cells are limited by the use of antigen-conjugated beads for stimulation and a relatively small number of identified pTyr sites, hindering comprehensive clinical applications.

In this study, we analyzed the pTyr proteome of clinical CAR-T products and captured approximately 40-170 pTyr sites. Most of these sites, especially those in the TCR signaling pathway proteins, were up-regulated, suggesting active tonic signaling. Notably, phosphorylation at JAK1_Y1034/1035 and STAT5_Y694 were significantly up-regulated, highlighting the activation of JAK-STAT signaling pathway, which is crucial for IL-2 induced cell proliferation and differentiation, suggesting high cell viability. We also noticed more acute pTyr changes in the TCR signaling pathway in CAR-T samples from the relapsed patients, potentially linked to the diminished therapeutic durability. Thus, controlling T cell activation during clinical CAR-T product preparation may be necessary.

To gain a comprehensive database of the tyrosine phosphorylation in CAR-T cells, a SILAC-labeled cell-cell co-culture assay using the antigen-presenting CD19-K562 cells to stimulate CAR-T cells was performed, with 766 pTyr sites identified. We analyzed the pTyr events under tonic and stimulated conditions, comparing two second generation CAR-T cells, i.e., 4-1BBζ and CD28ζ CAR-T cells, known for distinct signaling and clinical features. Both CAR-T types showed similar pTyr events, particularly in the TCR signaling pathway, with CD28 CAR-T cells exhibit higher changes, consisting with previous research.

Strikingly, comparison of clinical CAR-T data with our tyrosine phosphotproteome database revealed distinctive features of clinical CAR-T products, such as up-regulated pTyr sites in CARs, MAPKs (ERKs, JNKs and p38 proteins), PLCG1, DOK2, and HSCT, which showed significant up-regulation in primary CAR-T cells post-stimulation. In contrast, the pTyr sites in CD3E, PTPN11 and PAG1 showed opposite changes in these two datasets. Since those pTyr sites in MAPKs are positive regulators and CD3E and PTPN11 tyrosine phosporylation sites are negative regulators, these clinical CAR-T cells appeared to be in a hyper-activation state with activated feedback signaling pathway, potentially resulting in exhaustion or reduced efficacy post-infusion. Interestingly, inhibitors such as idelalisib, which target PIK3CD—a key regulator in the TCR signaling pathway—have been utilized during CAR-T cell manufacturing to enhance memory T cell formation^54–57^. Additionally, recent research has shown that MEK1/2 inhibitors, which target MAPK signaling pathway, can counteract CAR-T exhaustion and terminal differentiation^58^. Similarly, inhibitors targeting the significantly changed pTyr sites identified in this study could also improve the CAR-T manufacturing and clinical efficiency.

In summary, our results indicate that clinical CAR-T products may be in a hyper-activation state, with acute tyrosine phosphorylation changes in TCR pathway, potentially linked to CAR-T therapy relapse. Furthermore, we analyzed the tyrosine phosphorylation proteome of CAR-T cells under both tonic signaling and stimulated conditions, providing a comprehensive database for studying the CAR-T cell signal transduction and screening inhibitors to enhance in vivo CAR-T cell expansion and long-term persistence. Further larger clinical cohorts to confirm these conclusions are necessary due to the limited sample size in this study. pTyr sites identified in this study should also be verified to decipher the precise function in CAR-T signal transduction. Despite these limitations, our attempt offers novel insights into the quality control and efficacy prediction of clinical CAR-T products from the perspective of pTyr proteomics.

## Materials and Methods

### Clinical samples

Clinical samples were obtained from Shenzhen People’s Hospital and all procedures for sample collection have been ethically approved by the Medical Ethics Committee of the Shenzhen People’s Hospital, Shenzhen, China (LL-KY-2021479).

For collecting CAR-T cells from patient peripheral blood (PB) after CAR-T infusion, PB was centrifuged by density gradient sedimentation using Ficoll-Paque PLUS (GE Healthcare). PBMCs cells were then resuspended by auto MACS buffer and incubated with magnetic microbeads (Miltenyi Biotec) for collecting CD3+ T cells. MACS sorted T cells were then resuspended with staining buffer and further sorted and collected by flow cytometry.

### Cell lines

Lenti-X 293T cells (TaKaRa #632180) were grown in DMEM medium supplemented with 10% FBS, 100 U/mL penicillin, 100 mg/mL streptomycin at 37°C with 5% CO_2_. CD19-K562 stable cell line was established by transducing K562 myelogenous leukemia cells (ATCC #CCL-243) lentivirally with human CD19 gene, as described previously^59^. Established CD19-K562 cells were grown in RPMI 1640 supplemented with 10% FBS, 100 U/mL penicillin, 100 mg/mL streptomycin at 37°C with 5% CO_2_. All cells were tested and found to be free of mycoplasma contamination.

### CAR construction

The construction of CAR was conducted as described previously^59^. In brief, the CAR coding sequence was comprised by an anti-CD19 scFv fragment (FMC63), followed by a CD8a signal peptide, a CD8a hinge-transmembrane region and a human CD3z domain with the human 4-1BB or CD28 costimulatory domain. The EGFP coding sequence was also fused after an IRES sequence following the CAR sequence. Sequences coding CARs were then inserted into pHR-hEF1a vector via Gibson assembly cloning, respectively.

### Lentivirus preparation and infection

Lenti-X 293T cells were transfected with pCMVdR8.92 (Addgene #8455) and pMD2.G (Addgene #12259) and CAR-expression pHR-hEF1a plasmids via TranslT- LT1 transfection reagent (Mirus #2300). At 60 h.p.t., collect the media and centrifuge to remove cells. The viral media was aliquoted and stored at-80℃.

Human PBMCs were isolated from peripheral blood of healthy donors with Human T Cell Enrichment Kit (Stem Cell Technology #19051). Human T cells were cultured in RPMI-1640 medium supplemented with 10% FBS, 100 U/mL penicillin, 100 mg/mL streptomycin, and 100 U/mL recombinant hIL-2 (novoprotein). Before lentiviral infection, human T cells were activated by Human T-Activator CD3/CD28 Dynabeads (Life Technologies #11132D) at a bead-to-cell ratio of 3:1 for 24 h. Cells were then exposed to lentivirus for 18 h and Dynabeads were removed after 5 days. Medium and IL-2 were refreshed every 2-3 days^60^.

### Flow cytometry

MACS sorted CD3+ T cells were resuspended with staining buffer and incubated with anti-FMC63(PE) anti-body (FM3-HPY53-25test, ACROBiosystems) in the dark at room temperature for 15-20 minutes. After washed and resuspended, CAR-T cells were sorted and collected by BD FACSAria SORP flow cytometer (BD, Biosciences, USA).

### Cell-cell stimulation assay

The cell-cell coculturing stimulation assay was conducted from modified protocol as previously described^26^. CAR-T and CD19 cells were labeled in SILAC “light” (L-lysine and L-arginine) and “heavy” (L-[^13^C_6_, ^15^N_2_]-lysine and L-[^13^C_6_, ^15^N_4_]-arginine) separately. Before stimulation, starve CAR-T cell in FBS-free SILAC media for 1 h. 1 × 10^8^ CAR-T or CD19-K562 cells were washed and resuspended and then transferred onto ice for 10 min. CAR-T cells and CD19-K562 cells were then mixed together at a ratio of 1:1 with equal volume, quickly centrifuged at 2000 rpm for 5 min at 4 °C to promote cell–cell contact. Supernatants were then removed and cells were stimulated at 37 °C for 5 min without disrupting the cell pellet. After that, the cells were lysed in lysis buffer (50 mM Tris·HCl, pH 8.5, 8 M urea, 150 mM NaCl, 1% Triton X-100, 0.1% sodium deoxycholate, 1 mM sodium orthovanadate, protease inhibitors mixture (Complete mini; Roche), phosphatase inhibitor mixture (PhosSTOP; Roche)) immediately. For CAR-T cells stimulated at 0 min, the CAR-T and CD19-K562 cells were lysed separately and lysates were mixed together. Lysates were sonicated and centrifuged at 4 °C and supernatants were transferred to a new tube for precipitation by chloroform and methanol. Samples were then resuspended in urea buffer (8 M urea in 50 mM Tris-HCL, pH 8.5), reduced with DTT, alkylated with iodoacetamide, and digested with trypsin at 37 °C overnight. Trypsin digested peptides were desalted with SPE column, dried in speed vac and stored at-80 °C for further use.

### Enrichment of pTyr peptides

Peptides were resuspended in IAP buffer (50 mM MOPS, pH 7.2, 10 mM Na2HPO4, 50mM NaCl), and incubated with superbinder proteins^61^ purified by His beads at 4 °C overnight. Peptides were washed by IAP buffer supplemented with 20 mM imidazole and then eluted by elution buffer (500 mM imidazole in PBS). Peptides were further desalted by home-made C18 StageTip following by titanium dioxide (Ti^4+^-IMAC) enrichment as previously described^62,63^. Phosphor-peptides were then desalted again before LC-MS/MS analysis.

### LC-MS/MS

All data were collected using a Q-Exactive HF-X equipped with an Easy-nLC 1200 chromatography system (Thermo Fisher Scientific). The nanoLC separation was performed with an integrated spraytip column (20 cm × 100 μm i.d) packed with C4 (3 μm/120 Å, 0.5−0.8 cm, Dr. Maisch GmbH) and C18 (1.9 μm/120 Å, 19−20 cm, Dr. Maisch GmbH) at a flow rate of 250 nL/min. Peptides were separated with a binary buffer system of 0.1% FA (v/v) in water (buffer A) and 0.1% (v/v) FA in ACN (buffer B). Peptides were separated with an 80 min segmented gradient as follows: 4−8% (v/v) buffer B in 2 min, 8−28% (v/v) buffer B in 55 min, 28−40% (v/v) buffer B in 5 min, 40−97% (v/v) buffer B in 2 min, followed by a 16 min wash with 90% (v/v) buffer B. Full MS scans were acquired from m/z 350 to 1550 with a mass resolution of 120 000. The resolved fragments were scanned at mass resolution of 15 000 and AGC target value of 1E5. Top 30 parent ions with charge of 2 or higher were fragmented by higher energy collisional dissociation (HCD) fragmentation. Tyrosine phosphor-peptides were fragmented by HCD with normalized collision energy (NCE) of 25, respectively. The isolation window was 1.4 Da, and the dynamic exclusion time was 30 s with a max injection time of 50 ms.

### MS Data analysis

Raw files were searched using MaxQuant (version 1.6.14 and 2.1) with UniProtKB human proteome database (downloaded from UniProt on 27 March 2018) for database search. Oxidation (M), Deamidation (N) and Phospho (STY) were set as variable modifications while Carbamidomethyl (C) as a fixed modification. The LFQ (minimum ratio count 2, normalization type C) and the re-quantify was set with isobaric weight exponent 0.75. The PSM and protein FDR were set at 0.01 for identifications. As for label free quantification, the parameters that separate LFQ in parameter groups, stabilize large LFQ ratios and require MS/MS for LFQ comparisons were set as true. After search, data were further processed using Perseus (Version 2.0.6) and R software (Version 4.2.2). pTyr sites marked as reverse and potential contaminant were filtered out and localization probability > 0.75 were selected for further students t-test in Perseus. Processed data was then plotted in Prism (Version 10) or/and R with packages ggplot2 (Version 3.4.4), pheatmap (Version 1.0.12), Mfuzz (Version 2.56.0) and KSEA App^64^. For KEGG pathway and Gene Ontology enrichment, analyses were performed in Metascape^65^ and further plots were conducted in R. To investigate differences in the tyrosine phosphorylation proteome between relapsed and non-relapsed patients undergoing CAR-T therapy, a custom-developed “ClinicalCART” R package was integrated to facilitate the prediction of CAR-T treatment outcomes.

## Quantification and statistical analysis

Statistical significance was determined by two tailed Student’s t test. The LFQ intensities of pTyr sites were shown as mean ± SEM by three replicates.

## Data availability

Data reported in this study will be available upon request from the lead contact.

## Acknowledgements

The authors acknowledge funding from the Shenzhen Innovation of Science and Technology Commission (JCYJ20210324120210029, JSGGZD20220822095200001, JCYJ20200109141212325 and JCYJ20200109140814408), and Guangdong province (2019B151502050).

## Author Contributions

R.T. conceived the project, R.T., H.W., X.Z, and J.Z. designed the experiments. B.Y., X.Y., Q.K., W.C. and L.C. performed experiments. B.Y., X.Y., W.C. and A.H. analyzed data, W.L., X.W., and X.C. contributed in experiments and data analysis, C.F., G.L., L.H, X.Q., Y.C, and Y.L. contributed in clinical sample collection. B.Y., X.Y. and R.T. wrote the manuscript and all other authors edited the manuscript.

## Declaration of Interests

The authors declare no competing interests.

## Notes

### Competing Interest Statement

The authors have declared no competing interest.

